# Deep learning integration of molecular and interactome data for protein-compound interaction prediction

**DOI:** 10.1101/2021.01.31.429000

**Authors:** Narumi Watanabe, Yuuto Ohnuki, Yasubumi Sakakibara

**Author notes:** **Corresponding Author** Yasubumi Sakakibara, 3-14-1 Hiyoshi, Kohoku-ku, Yokohama, 223-8522, Japan.

## Abstract

**Motivation:** Virtual screening, which can computationally predict the presence or absence of protein-compound interactions, has attracted attention as a large-scale, low-cost, and short-term search method for seed compounds. Existing machine learning methods for predicting protein-compound interactions are largely divided into those based on molecular structure data and those based on network data. The former utilize information on proteins and compounds, such as amino acid sequences and chemical structures, while the latter utilize interaction network data, such as data on protein-protein interactions and compound-compound interactions. However, few attempts have been made to combine both types of data in molecular information and interaction networks.

**Results:** We developed a deep learning-based method that integrates protein features, compound features, and multiple types of interactome data to predict protein-compound interactions. We designed three benchmark datasets with different difficulties and evaluated the performance on them. The performance evaluations show that our deep learning framework for integrating molecular structure data and interactome data outperforms state-of-the-art machine learning methods for protein-compound interaction prediction tasks. The performance improvement is proven to be statistically significant by the Wilcoxon signed-rank test. This reveals that the multi-interactome captures different perspectives than amino acid sequence homology and chemical structure similarity, and both type of data have a synergistic effect in improving prediction accuracy. Furthermore, experiments on three benchmark datasets show that our method is more robust than existing methods in accurately predicting interactions between proteins and compounds that are unseen in the training samples.

## Introduction

Most compounds that currently act as drugs bind to target proteins that can cause disease, and these compounds can control their functions. Therefore, it is necessary to search for compounds that can interact with the target protein when developing new drugs, and this process must be performed efficiently. However, determining the interaction of a large number of protein-compound pairs via experiments is expensive in terms of time and cost.

Virtual screening that can computationally classify the presence or absence of protein-compound interactions has attracted attention as a large-scale, low-cost, short-term search method for hit compounds. In particular, the method of using machine learning for virtual screening is considered to be applicable to a wide variety of proteins and compounds.

Machine learning-based methods for predicting protein-compound interactions are largely divided into those based on molecular structure data and those based on network data. The former use protein and compound data represented in amino acid sequences and chemical structure formulas, and they can be applied to proteins when a docking simulation cannot be performed because the three-dimensional structure is unknown. In our previous study [1-3], using positive interactions between drug compounds and their target proteins downloaded from DrugBank (a database that contains information on existing drug compounds) [4] and negative interactions consisting of randomly combined compounds and proteins, we performed binary classification using a support vector machine (SVM). A prediction accuracy of 85.1% was achieved. Based on this result, we developed COPICAT, a comprehensive prediction system for protein-compound interactions, which enabled us to search for lead compounds from a huge compound database, PubChem [5], consisting of tens of millions of compounds.

Deep learning, a method developed in the field of machine learning, has been used in a variety of fields in recent years because it has achieved high prediction accuracy in fields such as image recognition, speech recognition, and compound activity prediction [6]. Deep learning-based protein-compound interaction prediction methods have been developed based on molecular structure data [7-10]. However, these existing deep learning-based methods utilize only information based on amino acid sequences and chemical structures, so the functional properties of proteins and compounds have not yet been incorporated into prediction.

The other type of machine learning approach for protein-compound interaction prediction is based on network data. An interaction network is commonly used to comprehensively represent interactions between molecules. For example, the protein-protein interaction network represents the relationships among physically interacting proteins. In the protein-protein interaction network, a node is a protein, and an edge is drawn between a pair of proteins that interact with each other.

Some previous studies incorporated data from multiple interaction networks to predict molecular interactions. For instance, multi-modal graphs were proposed to handle three types of interactions: protein-protein, protein-drug, and polypharmacy side effects. A deep learning method, Decagon [11], for multi-modal graphs was proposed to predict polypharmacy side effects. DTINet [12] and NeoDTI [13] were designed and developed as graph-based deep learning frameworks to integrate heterogeneous networks for drug-target interaction predictions and drug repositioning. In particular, NeoDTI exhibited a substantial performance improvement over other state-of-the-art prediction methods based on multiple interaction network data.

In addition to predicting protein-compound interactions, several studies have predicted other types of molecular interactions. Protein-protein interactions induce many biological processes within a cell, and experiential and computational methods have been developed to identify various protein-protein interactions. High-throughput experimental methods such as yeast two-hybrid screening were developed to discover and validate protein-protein interactions on a large scale. Computational methods for protein-protein interaction predictions employ various machine learning methods, such as SVM with feature extraction engineering [14]. The recurrent convolutional neural network (CNN), which is a deep learning method, was applied to sequence-based prediction for protein-protein interactions [15]. Compounds that can interact with each other are often represented as compound-compound interactions (also known as chemical-chemical interactions); interactive compounds tend to share similar functions. Compound-compound interactions, called drug-drug interactions, can be used to predict side effects based on the assumption that interacting compounds are more likely to have similar toxicity [16]. A computational approach to compound-compound interaction predictions has been studied with various machine learning methods, including end-to-end learning with a CNN based on the SMILES representation [17].

The purpose of this study is to improve prediction accuracy by integrating molecular structure data and heterogeneous interactome data into a deep learning method for predicting protein-compound interactions. In addition to the molecular information (amino acid sequence and chemical structure) itself, protein-protein interaction network data with similar reaction pathways or physical direct binding and compound network data linking compounds with similar molecular activities are incorporated into the deep learning model as multiple-interactome data. To the best of our knowledge, there are no deep learning-based solutions for predicting protein-compound interactions that integrate multiple heterogeneous interactome data along with the direct input of amino acid sequences and chemical structures.

This study proposes a method for predicting protein-compound (drug-target) interactions by combining protein features, compound features, and network context for both proteins and compounds. The network context comes in the form of protein-protein interactions from the STRING database [18], and the compound-compound interactions come from the STITCH database [19]. The protein-protein interaction network and compound-compound interaction network are processed using node2vec [20] to generate feature vectors for each protein node and each compound node in the interaction networks in an unsupervised manner. Each network-based representation is then combined with additional features extracted from a CNN applied to the amino acid sequence of a protein and from the extended-connectivity fingerprint (ECFP) of a compound. The final combined protein representations and compound representations are used to make a protein-compound interaction prediction with an additional fully connected layer. The overall learning architecture is illustrated in Figure 1.

**Figure 1.**
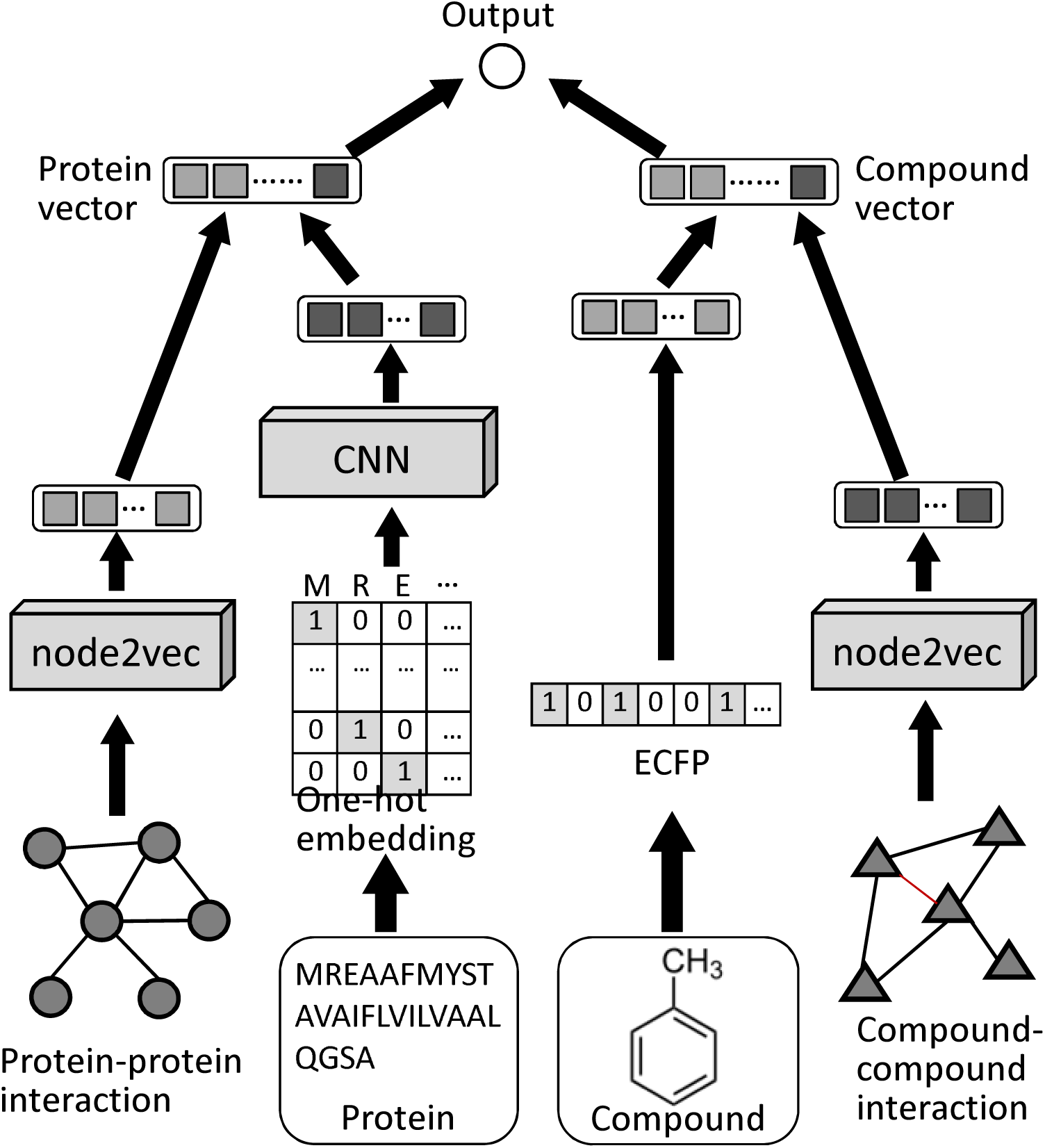
Deep learning architecture that integrates molecular structure data and interactome data to predict protein-compound interactions. It integrates graph-based and sequence-based representations for the target protein and compound. The amino acid sequence of the protein input was embedded into a one-hot vector of 20 dimensions in height. The ECFP representation of the compound input was embedded into a 1024-dimensional vector. The feature vectors were also extracted from the protein-protein and compound-compound interaction network using node2vec, a feature representation learning method for graphs. These feature vectors were combined as a protein vector and a compound vector. The interaction was predicted in the output unit.

We designed three benchmark datasets with different difficulties and evaluated the performance on them. In the performance evaluations, we demonstrate that integrating the molecular structure data and multiple heterogeneous interactome data has a synergistic effect in improving the accuracy of protein-compound interaction prediction. Furthermore, performance comparisons with state-of-the-art deep learning methods based on molecular information [10] and those based on interaction network data [13] as well as the traditional machine learning methods SVM and random forest show that our model exhibits significant performance improvements in the most important evaluation measures: AUROC, AUPRC, F-measure and accuracy, while the other methods show low values of these measures. The improvement is proven to be statistically significant by the Wilcoxon signed-rank test.

Finally, we analyse whether protein-protein interactions capture a different perspective than amino acid sequence homology and whether compound-compound interactions capture a different perspective than chemical structure similarity.

## Methods

### 1D-CNN for Encoding Protein Data

First, the protein data were applied to a one-dimensional convolutional neural network (1D-CNN). For the protein input, a one-hot vector was used for the distributed representation of an amino acid sequence of 20 dimensions at a height and width of 8,923 dimensions with the maximum length of amino acid sequences.

An amino acid sequence is a linear structure (1-D grid). In this study, a filter (kernel) with a one-dimensional convolution operation was applied to the linear structure. Here, a “one-dimensional” convolutional operation for linear structures was interpreted as scanning the input structure in only one direction along the linear structure with a filter of the same height (dimension) as that of the distributed representation of the input.

### One-Dimensional (1D) Convolutional Layer

We denote 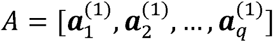 as an input vector sequence that corresponds to the one-hot vector representation of an amino acid sequence (as illustrated in Figure 1). For a filter function in the *l*-th hidden layer of the CNN, the input is the set of feature maps in the (*l-*1)-th hidden layer 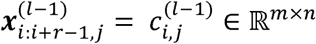, where *r* is the size of the filter, *m* is the size of the feature map, and *n* is the number of feature maps. The output of the *k*-th filter is a feature map of the *l*-th layer 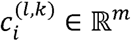, which is defined as follows:

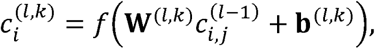

where *f* is an activation function (leaky-ReLU), **w**^*(l,k)*^∈ ℝ^*m×n×d*^is the weight matrix of the *k*-th filter in the *l*-th convolutional layer, and **b**^*(l,k)*^is the bias vector. The average-pooling mechanism is applied to every convolution output. To obtain the final output *y* = {*y*^(*t*,1)^, *y*^(*t*,2)^, …, *y*^(*t,s)*^ }, global max-pooling is used as follows:

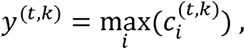

where *t* represents the last layer of the CNN and *s* represents the number of filters in the last layer.

### Extended-Connectivity Fingerprint (ECFP) for Compound Data

The extended-connectivity fingerprint (ECFP, also known as the circular fingerprint or Morgan fingerprint) [21] is the most commonly used feature representation for representing a property of the chemical structure of a compound. This algorithm first searches the partial structures around each atom recurrently, then assigns an integer identifier to each partial structure and expresses this as a binary vector by using a hash function. Potentially, an infinite number of structures exist in the chemical space; consequently, the ECFP requires vectors with a large number of bits (usually 1,024 - 2,048 bits). In this study, we employed an ECFP with 1024 bits as the feature representation for the chemical structure of a compound.

### Feature Representation Learning for Protein-protein and Compound-Compound Interactions

A protein-protein interaction network that connects physically interacting proteins and a compound-compound interaction network that connects compounds with similar molecular activities were input as multiple-interactome data. First, each network was represented as a graph. A node is a protein in the protein-protein network and a compound in the compound-compound network. An edge is drawn between a pair of proteins (compounds) that interact with each other. By applying this graph to “node2vec” [20], the feature vector of each node was obtained in an unsupervised manner; node2vec is a deep learning method that learns the feature representation of nodes in a graph and obtains a feature vector for each node.

Node2vec is a graph embedding algorithm that can be applied to any type of graph, and it can learn a feature vector such that nodes that are nearby on the graph are also close in the embedded feature space. In other words, the inner product of the feature vectors of the nearby nodes is high. It is known that the accuracy of the node classification task and the link prediction task using the obtained feature representations of nodes is higher than that of the existing methods.

The node2vec algorithm was applied to the protein-protein interaction network and the compound-compound interaction network. Using a protein and a compound as vertices, the interaction networks were converted into graphs with edge weights based on the reliability of the experimental data and the similarity in molecular activity. Node2vec (version 0.2.2) from the Python library, which implemented the node2vec algorithm, was applied to the converted graph. The node2vec parameters used the default values (embedding dimensions: 128; number of nodes searched in one random walk: walk_length=80; number of random walks per node: num_walk=10; control of probability of revisiting a walk node: p=1; control of the search speed and range: r =1; whether to reflect the graph weight: weight_key=weight).

Let a protein-protein interaction network be expressed by a weighted graph *G*_*protien*_*= (V*_*protien*_,*E*_*protien*_) and a compound-compound interaction network by a weighted graph *G*_*compound*_ *= (V*_*compound*_, *E*_*compound*_ ). By applying node2vec to these graphs, the feature representations can be obtained and are denoted as ***N***_*protien*_*=node2vec(Gprotien)* ∈ ℝ ^*d*^ *and ***N***_*compound*_ = node2vec(G*_*compound*_)∈ ℝ ^*d*^for a dimension of *d*.

### Deep Learning Model Structure for Integrating Molecular Information and the Interaction Network

The feature vectors obtained from the 1D-CNN for the amino acid sequence and node2vec for the protein-protein interaction network were concatenated and fed to the final output layer. Similarly, the feature vectors from the ECFP for the chemical structure and node2vec for the compound-compound interaction network were concatenated and fed to the final output layer.

We designed an output layer consisting of an element-wise product calculation followed by a fully connected layer, which is an extension of cosine similarity. The architecture is illustrated in Figure 2. First, the feature vectors for the proteins and compounds were mapped onto the same latent space with a fixed dimension *d* by applying fully connected layers. The similarity between the vector for proteins and the vector for compounds on the latent space was calculated by the element-wise product calculation method followed by a fully connected layer. When a pair of proteins and compounds was input, if the similarity was higher than some predefined threshold (where the default was 0.5), it was predicted that there was an interaction between the input pair. If the similarity was lower, it was predicted that there was no interaction. This model is denoted as the “integrated model”.

**Figure 2.**
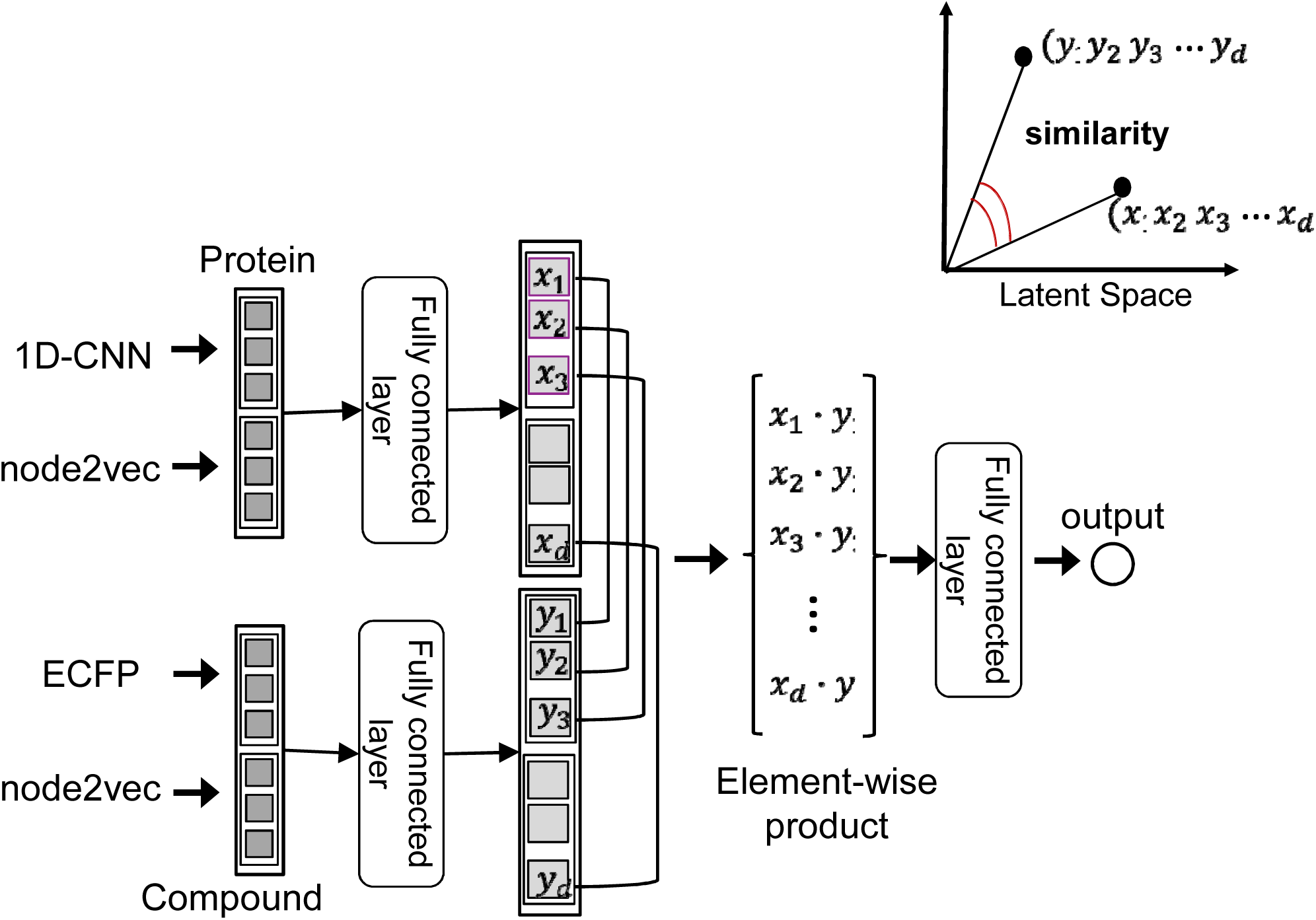
The output layer architecture. The integrated model predicts the protein-compound interactions by embedding the protein and compound data from different modalities into a common latent space. The feature vectors for the proteins and compounds are mapped onto the same latent space by applying a fully connected layer. Then, their similarity in the latent space is calculated with an element-wise product calculation followed by a fully connected layer.

More precisely, let ***a***_protien_denote the feature vector output by the 1D-CNN for an amino acid sequence, and let ***b***_compound_ denote the feature vector of the ECFP for the chemical structure of a compound. Let *N*_protien_and ***N***_compound_ denote the feature representations obtained from node2vec for the protein-protein interaction network and the compound-compound interaction network. Two feature vectors ***a***_protien_and ***N***_protien_were concatenated as one vector ***v***_protien_for the protein multi-modal feature. Two feature vectors ***b***_compound_ and ***N***_compound_ were concatenated as one vector ***v***_compound_ for the compound multi-modal feature. The concatenated feature vectors ***v***_protien_and ***v***_compound_ were mapped onto the same latent space with a fixed dimension *d* by applying the fully connected layers *f* and *g*. From this, the similarity between the two vectors for the latent space was calculated.

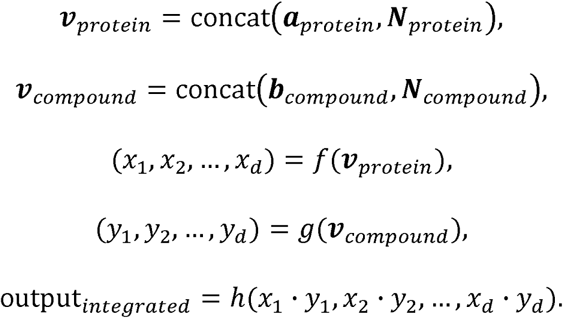

As described above, to handle data from different modalities such as proteins and compounds, we adopted a method of embedding data of different modalities into a common latent space. Defining the similarity in the obtained latent space enables the measurement of the similarity between the data for different modalities. Visual semantic embedding (VSE) is a typical example of a method that handles data from different modalities and can associate images with text data in acquiring these multi-modal representations [22]. VSE was developed to generate captions from images (image captioning). The image feature and the sentence feature are linearly transformed and embedded into a common latent space.

### Single-Modality Models

To see the effect of integrating multi-modal features, two baseline models were constructed for the performance comparison. One was based on molecular structure data and used only amino acid sequence and chemical structure information, and the other was based on interaction network data and used only protein-protein interaction and compound-compound interaction information. The single-modality model based on molecular structure data, denoted the “single-modality model (molecular)”, is defined as follows:

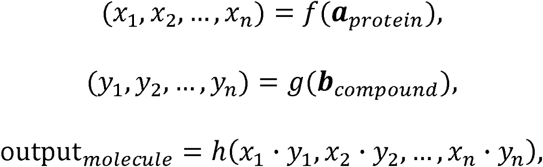

and the single-modality model based on interaction network data, denoted the “single-modality model (network)”, is defined as follows:

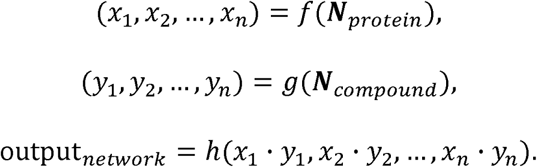

### Loss Function

For the similarity output *x* of the model, the output value was restricted to the range 0 to 1 by the sigmoid function, and cross entropy was applied as the loss function *L* (*θ*) tocalculate the training error.

### Hyperparameter Optimization

The hyperparameters, the number and size of the filters in the convolutional layers in the 1D-CNN, and the number of units in the fully connected output layers were optimized by the Bayesian optimization tool Optuna [23], which is an automatic hyperparameter optimization software framework particularly designed for machine learning. For the hyperparameter optimization, the validation dataset was obtained by dividing the training samples into a set for training and a set for validation.

### Regularization

Regularization is important for avoiding overfitting and improving the prediction accuracy in deep learning for complex model architectures with a large number of parameters.

Regularization is especially important in our deep learning model, which integrates multiple datasets of different modalities; hence, we employed several regularization methods.

We employed batch normalization [24], which allowed us to use much higher learning rates and be less careful about initialization, after each convolutional layer. We also inserted dropout [25] after the fully connected layers. Furthermore, we added an L2 regularization term to the training-loss function *L*(*θ*). When incorporating weight decay, the objective function to be optimized is as follows:

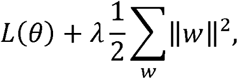

where *w* refers to the parameters of the entire model, and the second term of the above equation indicates taking the sum of the squared values of all the parameters and dividing by 2.λ is a parameter that controls the strength of regularization. Adding this term to the objective function has the effect of preventing the absolute value of the network weight from becoming too large, which helps prevent overfitting.

### Comparison with State-of-the-Art Existing Methods

The prediction performance of the proposed models was compared with that of state-of-the-art deep learning methods based on molecular structure data and interaction network data. The first method was based on a graph CNN for protein-compound prediction [10]. It employed a graph CNN for encoding chemical structures and a CNN for *n*-grams of amino acid sequences. The second method was NeoDTI [13], which demonstrated superior performance over other previous methods based on multiple-interaction-network data. We also compared our method with the traditional machine learning methods SVM and random forest [26] as the baseline prediction methods. These traditional methods require structured data as input. For the protein information, the 3-mer (3-residue) frequency in the amino acid sequence was used as the feature vector for 8,000 dimensions. For the compound information, an ECFP with a length of 1,024 and a radius of 2 was used. The radial basis function (RBF) was used as the kernel function of SVM, and all other parameters of SVM and random forest used the default values. In implementing these machine learning methods, sckit-learn (version 0.19.1) and chainer (version 5.0.0) were used.

### Datasets

The protein-compound interaction data and compound-compound networks were retrieved from the database STITCH [19], and the protein-protein networks were retrieved from the database STRING [18].

### Protein-Compound Interaction Data

Protein-compound interaction data can be obtained from the STITCH database [19]. STITCH contains data on the interaction of 430,000 compounds with 9.6 million proteins from 2,031 species. The STITCH data sources consist of (1) structure-based prediction results, such as the genome context and co-expression; (2) high-throughput experimental data; (3) automatic text mining; and (4) information from existing databases. When a protein-compound dataset is downloaded from STITCH, a score based on the reliability is created for each of the above four items for each protein-compound pair. For the protein-compound interaction data used in this study (as a “positive” example), the threshold value for the reliability score of item (2) was set to 700, and the data with a reliability score of 700 or higher were extracted from STITCH so that interologs were eliminated and the data were composed of only experimentally reliable interactions; the data that did not meet this threshold were removed.

For the STITCH data, interactions with a confidence score of 700 or more were determined based on the criterion that they were at least highly reliable [27]. Of the combinations of proteins and compounds, only pairs not stored in the STITCH database were taken as “negative” examples. In general, protein-compound pairs that are not stored in STITCH have very low confidence, with a score of 150 or less for their interaction [28], so these are considered to be non-interacting negative examples. The ratio of the positive and negative examples was 1 to 2.

### Protein-Protein Interaction Data

The protein-protein interaction information was obtained from the STRING database [18], which contains data for protein-protein interactions covering 24.6 million proteins from 5,090 species. The STRING data sources consist of (1) experimental data; (2) pathway databases; (3) automatic text mining; (4) co-expression information; (5) neighbouring gene information; (6) gene fusion information; and (7) co-occurrence-based information. In particular, item (1) is interaction data obtained from actual experiments, which include biochemical, biophysical, and genetic experiments. These are extracted from databases organized by the BioGRID database [29] and the IMEx consortium [30]. When the protein-protein interaction data from STRING were downloaded, a score based on the reliability was created for each of the above seven items for each protein-protein pair. Regarding the protein-protein interaction network, the threshold value for the reliability score of item (1) was set to 150. Data that did not satisfy this criterion were removed.

### Compound-Compound Interaction Data

The compound-compound interaction data were also obtained from the STITCH database. The compound-compound interaction data in STITCH are based on (1) the chemical reactions obtained from the pathway databases; (2) structural similarity; (3) association with previous literature; and (4) correspondence between the compounds based on molecular activity similarity. For the similarity of the molecular activities in item (4), the activity data obtained by screening the model cell line NCI60 were used. When the compound-compound interaction data were downloaded from STITCH, a score based on the reliability was created for each of the above four items for each compound pair. For the compound-compound interaction data used in this study, the threshold value for the reliability score in item (4) was set to 150. Data that did not satisfy this criterion were removed.

### Construction of the Baseline, Unseen Compound-Test, and Hard Datasets for Evaluation

From the STITCH and STRING databases, a total of 22,881 protein-compound interactions, 175,452 protein-protein interactions and 69,231 compound-compound interactions were downloaded. Using the downloaded dataset in which the protein-protein interaction, compound-compound interaction and protein-compound interaction data were all available, the three types of datasets below were constructed to perform five-fold cross validation. In typical *k*-fold cross validation, all positive and negative examples are randomly split into *k* folds. One of them is used as a test sample, and the remaining *k*−1 are used as training samples; then, the *k* results obtained are averaged. We call the cross-validation dataset the *baseline dataset*. In this study, as more difficult and more practical tasks, we constructed two more cross-validation datasets, called the *unseen compound-test dataset* and the *hard dataset*. In the unseen compound-test dataset, we split the data into *k* folds so that none of the folds contain the same compounds as the others. In the unseen compound-test dataset, the compounds in the test sample do not appear in the training sample. In other words, the interaction of new (unseen) candidate compounds with the target proteins must be accurately predicted. In the hard dataset, we split the data into *k* folds so that none of the folds contain the same proteins and compounds as the others. In the hard dataset, neither the proteins nor the compounds in the test sample appear in the training sample. In other words, interactions in which neither the proteins nor the compounds are found in the training sample must be accurately predicted.

## Results

The following measures were used for the performance evaluation criteria: AUROC (area under the receiver operating characteristic curve), AUPRC (area under the precision-recall curve), F-measure, and accuracy.

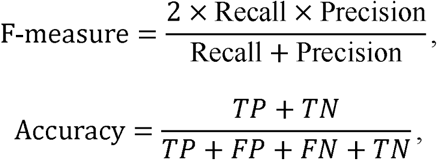

where *TP* is the number of true positives, *TN* is the number of true negatives, *FP* is the number of false positives, *FN* is the number of false negatives, Recall is defined by *TP*/(*TP*+*FN*), and Precision is defined by *TP*/(*TP*+*FP*).

### Effectiveness of Integrating Molecular Structure Data and Interaction Network Data

The performance of our three models was evaluated to determine the effectiveness of integrating the molecular structure data and the interaction network data. The results on the three datasets are shown in Tables 1-3. In the tables, the mean and standard deviation (SD) for the five folds are shown. Furthermore, the symbol “*” indicates that there was a significant difference in the Wilcoxon signed-rank test, with p-value p <0.05, in comparison with the integrated model.

**Table 1.** Performance comparison of three proposed models with existing methods on the baseline dataset.

|  | AUROC | AUPRC | F-measure | Accuracy |
| --- | --- | --- | --- | --- |
| Integrated model<br>(molecular+network) | **0.972±0.004** | **0.954±0.005** | **0.900±0.006** | **0.933±0.004** |
| Single-modality model<br>(molecular) | 0.956±0.004* | 0.927±0.006* | 0.868±0.009* | 0.911±0.006* |
| Single-modality model<br>(network) | 0.947±0.008* | 0.920±0.010* | 0.853±0.015* | 0.904±0.009* |
| Graph CNN<br>[10] | 0.917±0.006* | 0.850±0.006* | 0.794±0.014* | 0.864±0.008* |
| NeoDTI<br>[13] | 0.956±0.005* | 0.905±0.016* | 0.872±0.006* | 0.917±0.004* |
| SVM | 0.805±0.009* | 0.651±0.012* | 0.743±0.012* | 0.837±0.006* |
| Random forest | 0.873±0.009* | 0.767±0.015* | 0.837±0.012* | 0.895±0.007* |

**Table 2.** Performance comparison on the unseen compound-test dataset.

|  | AUROC | AUPRC | F-measure | Accuracy |
| --- | --- | --- | --- | --- |
| Integrated model<br>(molecular+network) | **0.890±0.039** | **0.842±0.050** | **0.727±0.085** | **0.843±0.038** |
| Single-modality model<br>(molecular) | 0.869±0.027 | 0.786±0.023* | 0.657±0.053 | 0.802±0.017 |
| Single-modality model<br>(network) | 0.831±0.053 | 0.759±0.055* | 0.661±0.073* | 0.809±0.030* |
| Graph CNN<br>[10] | 0.804±0.037* | 0.679±0.031* | 0.637±0.027 | 0.773±0.009* |
| NeoDTI<br>[13] | 0.823±0.067 | 0.773±0.064* | 0.621±0.062* | 0.805±0.024* |
| SVM | 0.765±0.020* | 0.603±0.029* | 0.689±0.029 | 0.810±0.016 |
| Random forest | 0.770±0.023* | 0.635±0.026* | 0.697±0.036 | 0.828±0.014 |

**Table 3.**
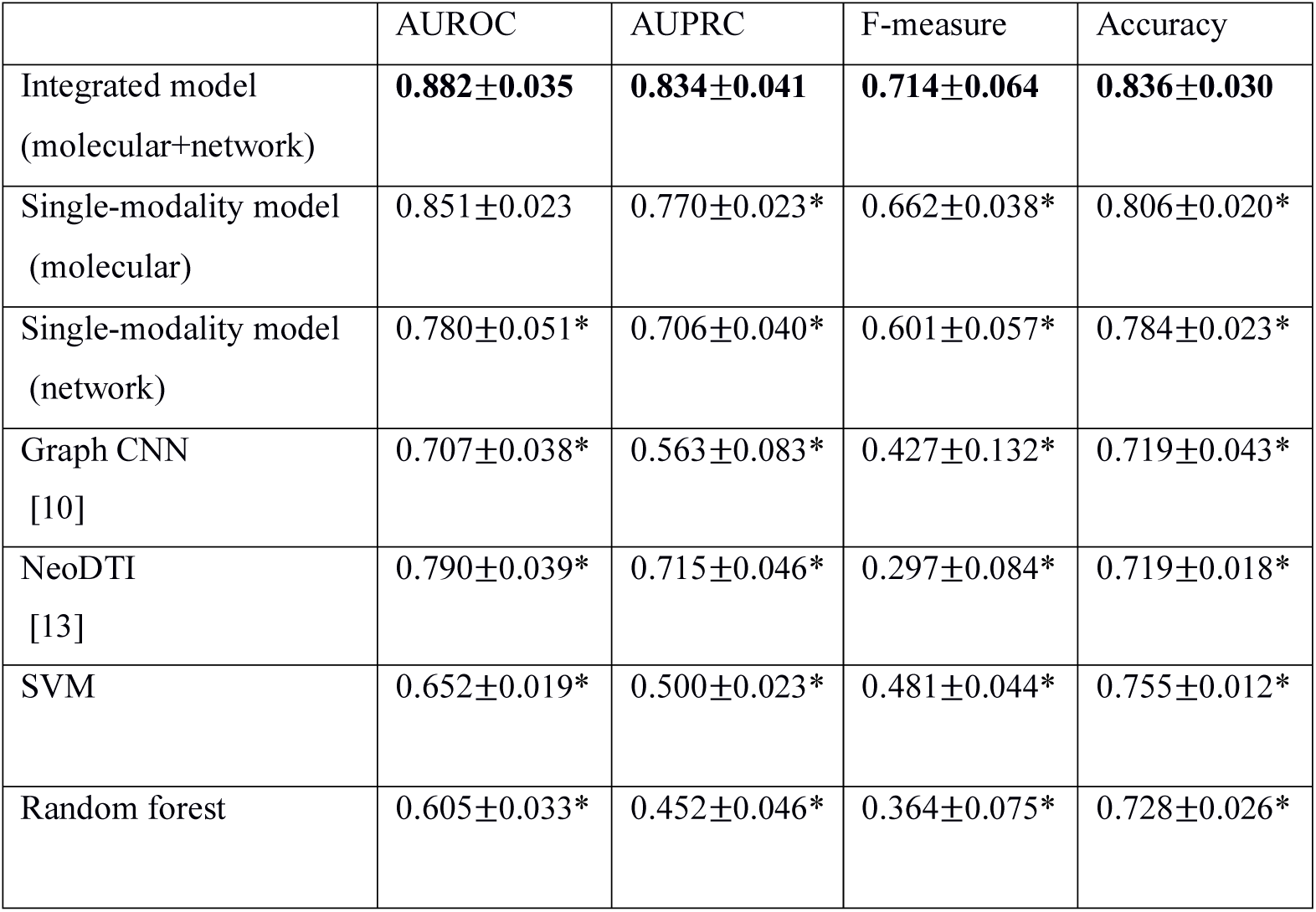
Performance comparison on the hard dataset.

Compared with the two single-modality models, the integrated model significantly improved the prediction accuracy in all evaluation measures. For example, in terms of AUPRC, which is a more informative evaluation index in a dataset that is imbalanced between positive and negative samples, the integrated model showed significant improvements of 3.0%, 7.1% and 8.3% over the single-modality model (molecular) and 3.7%, 10.9% and 18.1% over the single-modality model (network) in the baseline dataset, the unseen compound-test dataset and the hard dataset, respectively. This demonstrates that integrating multiple heterogeneous interactome data with molecular structure data brought a synergistic effect in improving the accuracy of protein-compound interaction prediction.

### Performance Comparison with Other Existing Methods

The prediction performance of our three models was compared with that of state-of-the-art deep learning methods and traditional machine learning methods based on molecular structure data and interaction network data. The results on the three datasets are shown in Tables 1-3.

The integrated model yielded superior prediction performance compared with the other existing methods. In the baseline dataset, the integrated model achieved significant improvements compared with the graph CNN-based method [10], NeoDTI [13] and the traditional machine learning methods SVM and random forest (Table 1). In fact, the Wilcoxon signed-rank test [31] verification showed that the performance difference was statistically significant, with a p-value p<0.05, and hence proved the superiority of the integrated model.

In the unseen compound-test dataset and the hard dataset, a more remarkable difference in the performance of the integrated model was confirmed. We compared the integrated model with the graph CNN-based method and NeoDTI in terms of AUROC, AUPRC and F-measure. The integrated model greatly outperformed the others, with significant improvements (10.7% in terms of AUROC, 24.0% in terms of AUPRC and 14.1% in terms of F-measure on the unseen compound-test dataset, and 24.8% in terms of AUROC, 48.1% in terms of AUPRC and 67.2% in terms of F-measure on the hard dataset) over the graph CNN-based method. In comparison with NeoDTI, significant improvements were also confirmed: 8.1% in terms of AUROC, 8.9% in terms of AUPRC and 17.1% in terms of F-measure on the unseen compound-test dataset, and 11.6% in terms of AUROC, 16.6% in terms of AUPRC and 140.4% in terms of F-measure on the hard dataset. Based on the above results, the integrated model can predict protein-compound interactions with stable accuracy, regardless of the difficulty of the dataset and the types of proteins and compounds that make up the test data, compared to other existing methods. This is due to the integrated model using features based on sequence information and compound structure information and features obtained from the interaction network as well as the effect of using the element-wise product of the protein feature vector and the compound feature vector in the output layer.

The single-modality model also yielded superior prediction performance compared with the existing methods using the same-modality input data. The graph CNN-based prediction method [10] obtains a compound feature vector by converting the chemical structure into a graph and applying it to the graph CNN, and it obtains a protein feature vector by splitting the amino acid sequence into *n*-grams and applying it to the CNN. Therefore, the graph CNN-based method can be defined as having the same molecular structure data-based prediction model as the single-modality model (molecular). In the baseline dataset, the unseen compound-test dataset and the hard dataset, the single-modality model (molecular) outperformed the graph CNN-based prediction method. For example, in the hard dataset, the single-modality model (molecular) achieved an improvement of 20.4% in terms of AUROC, 36.8% in terms of AUPRC and 55.0% in terms of F-measure on the hard dataset over the graph CNN-based method (Table 3). From this result, in protein-compound interaction prediction, it is sufficient to use the ECFP as a feature representation for the compound structure, compared with the deep learning method in which the compound structure is converted into a graph structure and a graph CNN is applied.

NeoDTI takes protein-protein interaction and compound-compound interaction information as input and predicts whether an edge is drawn between the compound and protein nodes by learning to reconstruct the network. Therefore, NeoDTI can be defined as an interaction network-based prediction model, which is the same as the single-modality model (network). The difference is that the single-modality model (network) first uses unsupervised deep learning (node2vec) to automatically learn feature representations for nodes in the given heterogeneous interaction networks and then applies supervised learning to predict protein-compound interactions based on the learned features, while NetoDTI simultaneously learns the feature representations of nodes and protein-compound interactions in a supervised manner. In the three datasets, the prediction performance of the single-modality model (network) was comparable to that of NetoDTI.

## Discussion

To interpret the accuracy improvement obtained by integrating multiple interactome data with molecular structure data, which was shown in the previous section, we analysed whether the protein-protein interaction captured a different perspective than amino acid sequence homology and whether the compound-compound interaction captured a different perspective than chemical structure similarity. More concretely, we investigated the relationship between the amino acid sequence homology and the similarity of proteins in the protein-protein interaction network as well as the relationship between the chemical structure similarity and the similarity in the compound-compound interaction network.

For every pair of proteins in the dataset used in the experiments, the amino acid sequence similarity was calculated using DIAMOND, and the cosine similarity between two vectors of the pair output by node2vec using the protein-protein interaction network was calculated. All of the protein pairs were plotted with the amino acid sequence similarity on the x-axis and the cosine similarity in the protein-protein interaction network on the y-axis. The scatter plot is shown in Figure 3 (top). Similarly, for every pair of compounds, the Jaccard coefficient of the ECFPs of the two compounds and the cosine similarity between the two vectors output by node2vec using a compound-compound interaction network were calculated. All of the compound pairs were plotted with the Jaccard coefficient on the x-axis and the cosine similarity in the compound-compound interaction network on the y-axis, as shown in Figure 3 (bottom). In both scatter plots, no clear correlation was observed. In fact, the correlation coefficients for each scatter plot were 0.186 and 0.199, respectively. In other words, it was confirmed that the amino acid sequence similarity and the similarity in the protein-protein interaction network were not proportional. Similarly, it was confirmed that the chemical structure similarity and the similarity in the compound-compound interaction network were not proportional. Therefore, we concluded that the protein-protein interaction network captured a different perspective than the amino acid sequence homology and compensated for it. The compound-compound interactions captured a different perspective than the chemical structure similarity and compensated for it.

**Figure 3.**
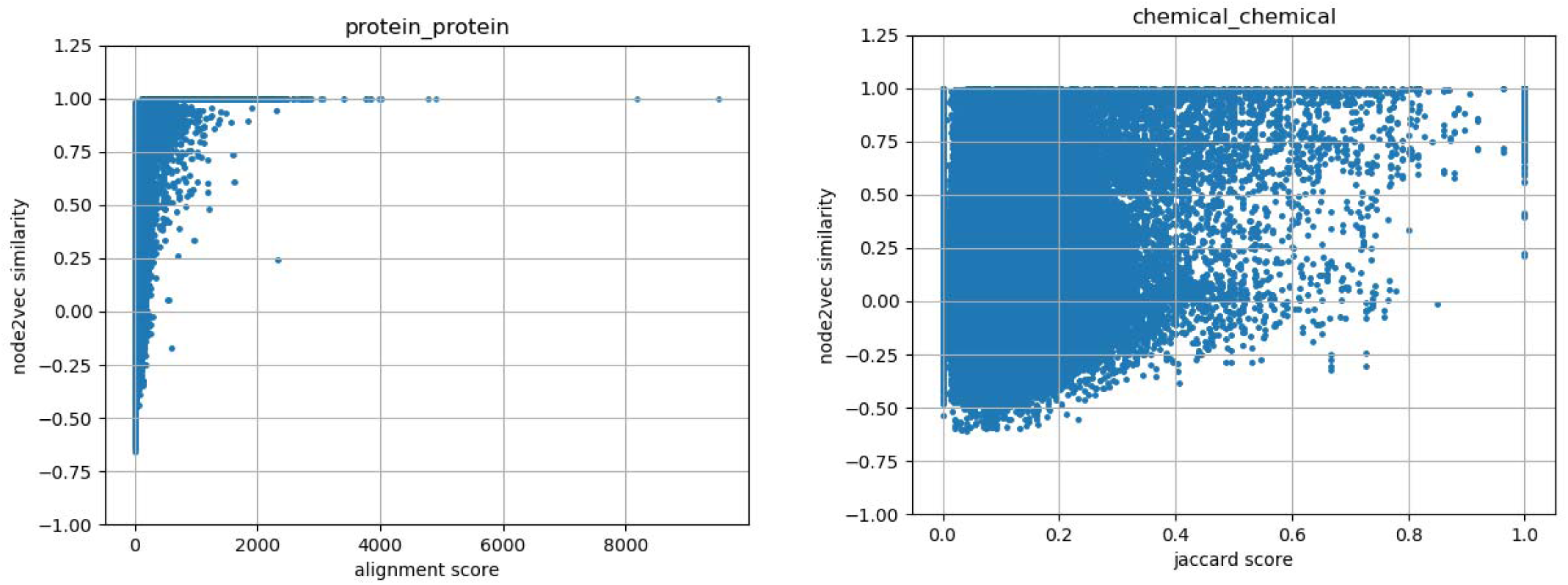
(Top) Relationship between the amino acid sequence similarity and the similarity in protein-protein interaction network. (Bottom) Relationship between the chemical-structure similarity and the similarity in compound-compound interaction network. The amino acid sequence similarity was calculated using DIAMOND, and the chemical structure similarity was calculated as the Jaccard coefficient of the ECFPs of the two compounds. The correlation coefficients are 0.186 and 0.199, respectively.

For example, the protein “5-hydroxytryptamine (serotonin) receptor 6, G protein-coupled (HTR6)” and the compound “Mesulergine” in the test sample in the “hard dataset” have a positive interaction [32], and our model succeeded in correctly predicting it. However, the single-modality model (molecular) and graph CNN-based method failed to predict the positive interaction; that is, both predicted that the pair would not interact. The most similar protein-compound pair in the training sample to the pair HTR6 and Mesulergine was the protein “adrenoceptor alpha 2A (ADRA2A)” and the compound “Pergolide” [33]. The protein ADRA2A and the compound Pergolide have a positive interaction in the training sample. The sequence similarity score between HTR6 and ADRA2A is rather low at 100.5, but the similarity of the two proteins in the protein-protein interaction network is relatively high at 0.805. A part of the protein-protein interaction network around HTR6 and ADRA2A is displayed in Figure 4 (left). Similarly, the Jaccard coefficient of the ECFPs between Mesulergine and Pergolide is relatively low 0.273 (in general, compound pairs with a Jaccard coefficient of ECFPs below 0.25 are considered not to have chemically similar structures [34]), but the cosine similarity of the two compounds in the compound-compound interaction network is high at 0.735. A part of the compound-compound interaction network around Mesulergine and Pergolide is displayed in Figure 4 (right).

**Figure 4.**
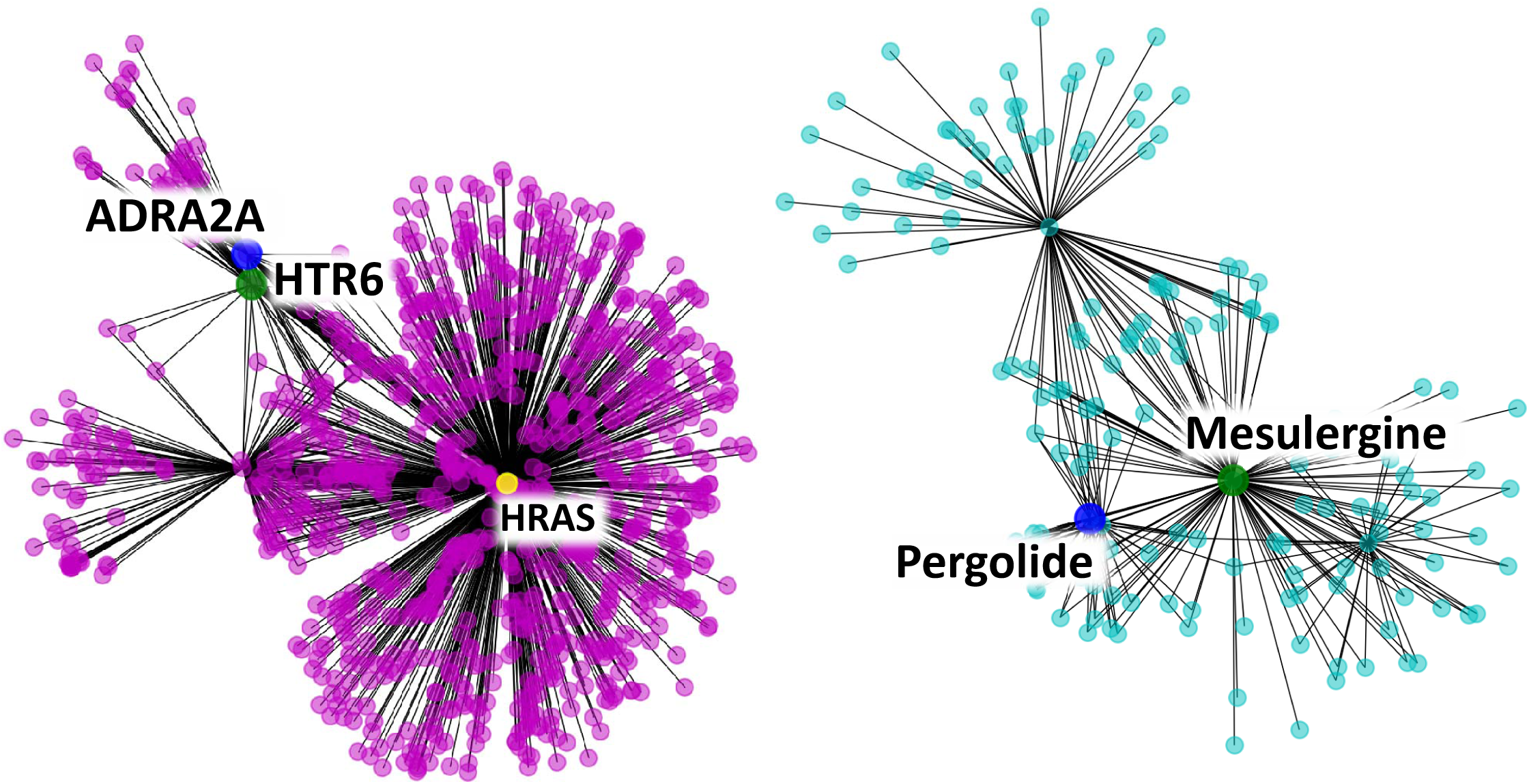
(Left) Part of the protein-protein interaction network around ABL1 and YES1. (Right) Part of the compound-compound interaction network around Crizotinib and Ceritinib.

## Conclusions

This study aimed to improve the performance of predicting protein-compound interactions by integrating molecular structure data and interactome data. This was achieved by integrating multiple heterogeneous interactome data into predictions of protein-compound interactions. An end-to-end learning method was developed that combined a 1D-CNN for amino acid sequences, an ECFP representation for compounds, and feature representation learning with node2vec for protein-protein and compound-compound interaction networks. The proposed integrated model exhibited significant performance differences with respect to the accuracy measures in comparison to the current state-of-the-art deep learning methods. The performance improvement was verified by the Wilcoxon signed-rank test as being statistically significant. The results indicated that the proposed model was able to more accurately predict the protein-compound interactions even in the hard dataset, where neither the proteins nor the compounds in the test sample appear in the training sample.

An important future task is to integrate the gene regulatory network as additional interactome data to further improve protein-compound interaction prediction. A large number of gene expression profiles for various tissues and cell lines are available in public databases, and gene regulatory networks can be effectively inferred from the gene expression profiles.

## List of Abbreviations

SVM: Support Vector Machines
CNN: Convolutional Neural Network
ECFP: Extended-Connectivity Fingerprint
VSE: Visual Semantic Embedding
AUROC: Area Under the Receiver Operating characteristic Curve
AUPRC: Area Under the Precision-Recall Curve
SD: Standard Deviation

## Declarations

### Availability of Data and Materials

The source code for the implementation of this deep learning method, along with the dataset for the performance evaluation, is available at https://github.com/Njk-901aru/multi_DTI.git.

## Authors’ Contributions

NW; Implemented the software, analysed data, and co-wrote the paper. YO; analysed data and compared with the existing methods. YS; designed and supervised the research, analysed data, and co-wrote the paper. All authors read and approved the final manuscript.

## Acknowledgements

Not applicable.

